# Fine-scale landscape genetics unveiling contemporary asymmetric movement of red panda (*Ailurus fulgens*) in Kangchenjunga landscape, India

**DOI:** 10.1101/2020.08.26.268532

**Authors:** Supriyo Dalui, Hiren Khatri, Sujeet Kumar Singh, Shambadeb Basu, Avijit Ghosh, Tanoy Mukherjee, Lalit Kumar Sharma, Randeep Singh, Kailash Chandra, Mukesh Thakur

## Abstract

Wildlife management in rapid changing landscapes requires critical planning through cross cutting networks, and understanding of landscape features, often affected by the anthropogenic activities. The present study demonstrates fine-scale spatial patterns of genetic variation and contemporary gene flow of red panda (*Ailurus fulgens*) populations with respect to landscape connectivity in Kangchenjunga Landscape (KL), India. The study found about 1309.54 Km^2^ area suitable for red panda in KL-India, of which 62.21% area fell under the Protected Area network. We identified 24 unique individuals from 234 feces collected at nine microsatellite loci. The spatially explicit and non-explicit Bayesian clustering algorithms evident to exhibit population structuring and supported red panda populations to exist in meta-population frame work. In concurrence to the habitat suitability and landscape connectivity models, gene flow results supported a contemporary asymmetric movement of red panda by connecting KL- India in a crescent arc. We demonstrate the structural-operational connectivity of corridors in KL-India that facilitated red panda movement in the past. We also seek for cooperation in Nepal, Bhutan and China to aid in preparing for a comprehensive monitoring plan for the long-term conservation and management of red panda in trans-boundary landscapes.

## Introduction

Habitat mapping and modelling corridors across species distribution are cardinal for prioritization of conservation strategies [1, 2]. Landscape connectivity demonstrates feasibility for wildlife to move through fragmented habitats and therefore maintaining corridors in fragmented landscapes are vital to ensure natural gene flow and the long-term survival of the species [1, 3]. Further, heterogeneity and rapid changes imposed in the landscape often accelerate restriction in the species movement between suitable patches [4, 5]. This restricted movement may lead to genetic consequences including disruption of gene flow, inflation of inbreeding and loss of rare alleles supporting local adaptation and genetic fitness [6, 7]. This phenomenon may induce multifaceted challenges in small populations inhabiting in trans-boundary landscapes (TBL). Prioritizing species conservation across TBL is challenging due to differences in the national interest, policies, local communities, funds allocation and political will [8].

The Kangchenjunga Landscape (KL) is one of the six TBL in the Hindu Kush Himalayan region, sharing boundary among Nepal, India and Bhutan [9]. The red panda (*Ailurus fulgens*), a magnificent iconic species of this landscape, is endemic to temperate conifer and cool broadleaf forest with dense bamboo undergrowth of preferring altitude range 2300 to 4000 m of Central and Eastern Himalayan biotic province [10–12]. Red Panda was taxonomically classified to occur in two subspecies based on the morphology and distribution - *Ailurus fulgens fulgens* distributed in Nepal, India, Bhutan, Myanmar, and China (Tibet and western Yunnan province) and *Ailurus fulgens styani*, occurred from the Sichuan and Yunnan provinces of China and Nujiang river was believed to be a biogeographic barrier for the separation of two subspecies [13,14]. However, a recent study by Hu et al. [15] demonstrated the presence of two distinct species of red panda, *i.e.* the Himalayan red panda (*Ailurus fulgens*) and Chinese red panda (*Ailurus styani*), with high depth sequencing data and revealed that the Yalu Zangbu river might be the potential boundary for species distribution. Further, Himalayan red panda reported to be distributed in India, Nepal, Bhutan, northern Myanmar, and Tibet and western Yunnan Province of China, while the Chinese red panda distributed in the Yunnan and Sichuan provinces of China [15]. The anthropogenic factors including habitat loss, poaching for pelt, jhoom cultivation and conversion of forest to non-forest land use have caused rapid decline of red panda [12,14,16]. Nearly 50% of red panda habitat has been lost in the last three generations, bringing it as ‘*Endangered’* in the Red list category [12]. In India, the red panda is protected under *Schedule-I* of the Wildlife (Protection) Act, 1972 and most of its populations reside in small and isolated protected areas (<500 sq. km), thereby increasing high risk of local extirpation of red pandas due to genetic inbreeding and loss of heterozygosity [15,17,18,19]. Earlier studies available on red panda from KL-India have addressed population status, distribution, and abundance [11, 20, 21], habitat preferences and diet composition [22, 23]. However, with the emergence of landscape genetics, it is now feasible to explicitly quantify the effects of landscape features on the spatial patterns of genetic variation, population structure, gene flow, and adaptation [24–26]. Thus, population genetics integrated with landscape ecology and remote sensing data can be used to aid delineating shift, if any, in the identified corridors that maintain connectivity between habitat patches and facilitate biotic processes such as dispersal and gene flow [27–30]. In this view, the detailed population genetic assessment of red panda with respect to landscape connectivity and anthropogenic activities is imperative to prioritize the management strategies for ensuring long term population viability of red panda in Himalayas. The present study is aimed to address the fine-scale spatial patterns of genetic differentiation and gene flow among the habitat clusters supporting red panda population in KL-India

## Results

Forty-eight candidate maxent models were generated (Table S3) and model with lowest Delta AICc informative AUC (0.911 ±0.098) was selected, which predicted species distribution better than the random model (Table S1). The generated binary map discriminated between suitable and unsuitable habitat of red panda based on 10-percentile training presence thresholds (Fig 1b). The jackknife test revealed that bioclimatic variable siwb_bio19, precipitation of coldest quarter contributed 45.6% and maximum temperature of the warmest month contributed 29.4% to predict habitat suitability. The land use land cover (LULC) contributed 12.9% followed by canopy height 11.8% and isothermality found to be less important and contributed only 0.3% (table 2). The Jackknife test for regularized training gain in the present model showed that the variable with the highest gain, when used in isolation, was precipitation of coldest quarter, which therefore appeared to have the most useful information by itself. The environmental variable that decreased the gain the most when it was omitted, was Max Temperature of Warmest Month, which therefore appeared to have the most information that was not present in the other variables (Fig S1).

**Fig 1.**
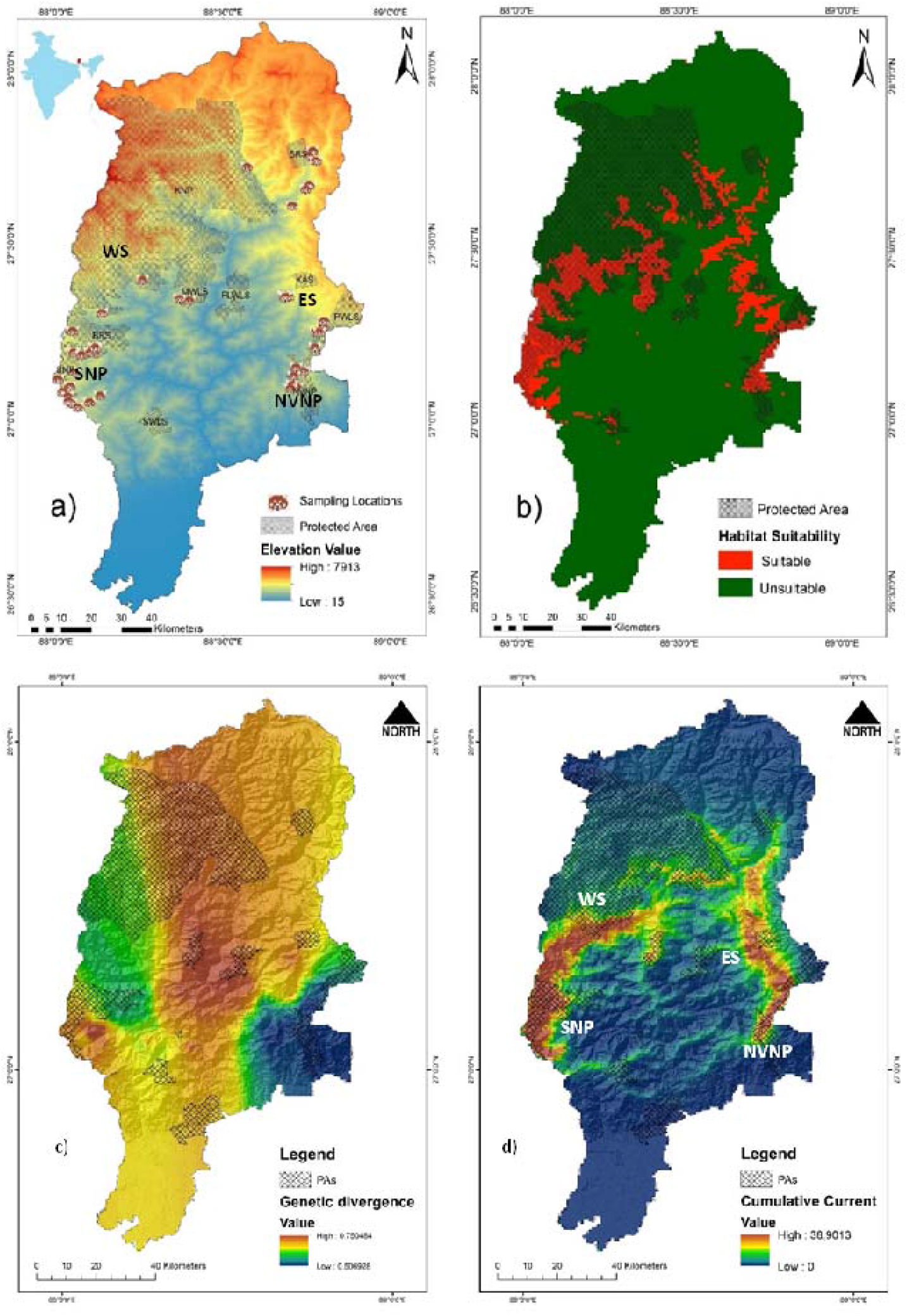
Study area map, red panda species distribution and landscape connectivity model. a) Map of Kangchenjunga landscape (KL), India with overlaid sampling locations [SNP-Singalila National Park and NVNP-Neora Valley National Park in north West Bengal; BRS-Barsey Rhododendron Wildlife Sanctuary and KNP-Kanchenjunga National Park in West Sikkim (WS); PWLS-Pangolakha Wildlife Sanctuary and KAS-Kyongnosola Alpine sanctuary in the East Sikkim (ES)], b) Predicted habitat suitability model of red panda in KL-India, c)Model based on genetic divergence of red panda in KL-India. d) Landscape connectivity model based on ensemble approach by combining both the genetic divergence and environmental conductance in KL-India.

### Habitat suitability and landscape connectivity

The study covered 10 protected areas (PAs) that included Singalila National Park (SNP), Senchal Wildlife Sanctuary (SWLS), and Neora valley National Park (NVNP) located in Northern West Bengal and Barsey Rhododendron Sanctuary (BRS), Maenam Wildlife Sanctuary (MWLS), FambongLho Wildlife Sanctuary (FLWLS), Pangolakha Wildlife Sanctuary (PWLS), Kangchenjunga National Park (KNP), Shingba Rhododendron Sanctuary (SRS), and Kyongnosla Alpine Sanctuary (KAS) located in Sikkim State of India. Our results predicted a total of 1309.54 Km^2^ (12.78%), comprised of 1097.26 Km^2^ in Sikkim and 212.28 Km^2^ in North West Bengal as suitable habitat for red panda with largest suitable habitats in KNP (493.37 Km^2^) (Table 1). The habitat suitability demonstrated that from West to East, SNP was connected with BRS and BRS was connected with KNP which is the only National Park in Sikkim. The MWLS in south Sikkim was connected with KNP on its northern border. On eastern part of the landscape NVNP and PWLS were connected, however, large amount of habitat suitability was observed outside PA on the eastern part of Sikkim.

**Table 1.**
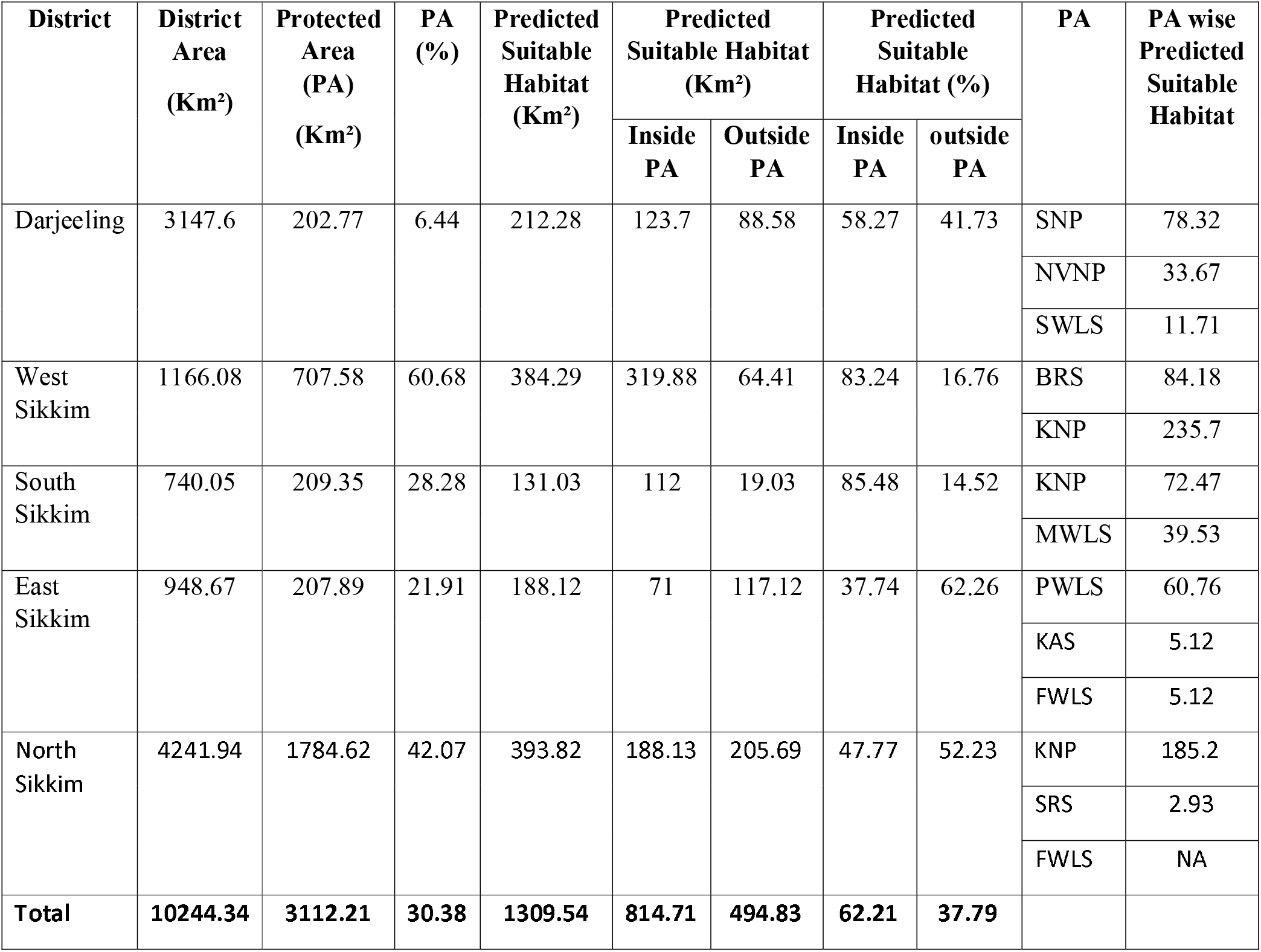
District and protected area wise habitat suitability

| District | District Area (Km<sup>2</sup>) | Protected Area (PA) (Km<sup>2</sup>) | PA (%) | Predicted Suitable Habitat (Km<sup>2</sup>) | Predicted Suitable Habitat (Km<sup>2</sup>) |  | Predicted Suitable Habitat (%) |  | PA | PA wise Predicted Suitable Habitat |
| --- | --- | --- | --- | --- | --- | --- | --- | --- | --- | --- |
|  |  |  |  |  | Inside PA | Outside PA | Inside PA | outside PA |  |  |
| Darjeeling | 3147.6 | 202.77 | 6.44 | 212.28 | 123.7 | 88.58 | 58.27 | 41.73 | SNP | 78.32 |
|  |  |  |  |  |  |  |  |  | NVNP | 33.67 |
|  |  |  |  |  |  |  |  |  | SWLS | 11.71 |
| West Sikkim | 1166.08 | 707.58 | 60.68 | 384.29 | 319.88 | 64.41 | 83.24 | 16.76 | BRS | 84.18 |
|  |  |  |  |  |  |  |  |  | KNP | 235.7 |
| South Sikkim | 740.05 | 209.35 | 28.28 | 131.03 | 112 | 19.03 | 85.48 | 14.52 | KNP | 72.47 |
|  |  |  |  |  |  |  |  |  | MWLS | 39.53 |
| East Sikkim | 948.67 | 207.89 | 21.91 | 188.12 | 71 | 117.12 | 37.74 | 62.26 | PWLS | 60.76 |
|  |  |  |  |  |  |  |  |  | KAS | 5.12 |
|  |  |  |  |  |  |  |  |  | FWLS | 5.12 |
| North Sikkim | 4241.94 | 1784.62 | 42.07 | 393.82 | 188.13 | 205.69 | 47.77 | 52.23 | KNP | 185.2 |
|  |  |  |  |  |  |  |  |  | SRS | 2.93 |
|  |  |  |  |  |  |  |  |  | FWLS | NA |
| **Total** | **10244.34** | **3112.21** | **30.38** | **1309.54** | **814.71** | **494.83** | **62.21** | **37.79** |  |  |

The ensemble approach by combining both the genetic divergence and environmental conductance depicted the landscape connectivity and its functionality to support movement corridors of red panda in KL-India. The predicted genetic divergence among the identified clusters was in coherence to the habitat suitability model (Fig 1b and 1c). The landscape genetics model by an ensemble conductance surface, lumping both the genetic divergence and environmental suitability demonstrated high-density currents radiated from SNP and NVNP in the North West Bengal (Fig.1d). Further, to decipher the direction of gene flow among the clusters, we estimated the contemporary migration rate using BayesAss (Fig S4). The landscape connectivity model and the contemporary migration rate, jointly inferred the permeability of floating currents, indicating structural connectivity in KL- India (Fig 1d), well supported by the asymmetric gene flow from West to East (Fig S4). The landscape connectivity model did not show horizontal connection from West to East in KL- India. Instead, the current flow forming a crescent arc to connect KL -India from the West to East with high-density currents radiating from SNP and NVNP of North West Bengal. The current formed bottleneck position of corridor in the KNP of north Sikkim. On the eastern part of the KL, the cumulative current flow was narrow running in longitudinal extent between PAs (Fig 1d).

### Individual identification and assessment of genetic variability

Allelic Drop out (ADO) ranged from 0 to 0.294 while we did not observe any significant amount of False allele (FA) (Table 3). Four loci i.e. CRP357, CRP385, CRP409 and CRP367 exhibited relatively high frequencies of null alleles that may consequently affected the Hardy Weinberg Equilibrium (HWE) and Inbreeding coefficient (F_IS_) (Table 3). Among 234 faecel samples collected from KL, we obtained positive PCR amplification in 98 faeces (38%) with nine microsatellite loci. Of which, 37 samples (approx. 40%) yielded mixed profiles on capillary electrophoresis, plausibly due to the red panda behaviour, being arboreal and group defecation in piles at same places. We finally obtained 57 unambiguous genotypes and with a select panel of seven loci with a cumulative P_IDsib_ 2.91×10^−3^, we identified 24 unique genotypes-two individuals originated from SNP, 11 from WS, six from ES and five from NVNP (Table 3). We uploaded multilocus genotype data of 24 unique individuals on DRYAD and available on https://doi.org/10.5061/dryad.2280gb5nz. For population genetic analysis, we used nine microsatellite data that exhibited ≥90 % amplification success. The exhibited mean observed number of alleles (Na)- 6±0.58, observed heterozygosity (H_O_)- 0.493±0.06 and expected heterozygosity (H_E_) 0.664±0.027 (Table 3). Three loci, CRP357, CRP385 and CRP367 deviated significantly from HWE and only one pair of loci out of 36 pair-wise comparisons were in significant Linkage disequilibrium (LD) (P<0.0012) after Bonferroni correction. HWE deviation of these markers might be due to the multiple reasons, like inflated frequencies of null alleles, Wahlund effect, and inbreeding due to consanguineous and assortative mating [31]. The mean F_IS_ was 0.276, indicating a significant inbreeding and loss of heterozygosity (Table 3).

**Table 2.** Details of bio-climatic variables used in the present study to predicted habitat suitability

| Variable | Code | Contribution (%) |
| --- | --- | --- |
| Isothermality (BIO2/BIO7) (* 100) | siwb_bio3 | 0.3 |
| Max Temperature of Warmest Month | siwb_bio5 | 29.4 |
| Precipitation of Coldest Quarter | siwb_bio19 | 45.6 |
| Land Use Land Cover | siwb_lulc_1k | 12.9 |
| Canopy Height | siwb_canopy_height | 11.8 |

**Table 3.** Genetic diversity indices and genotyping error in red panda population at nine microsatellite loci

| Locus | Na | Ho | He | Fis(W&C) | PID (locus) | PIDsib (locus) | F<sub>Null</sub> | Allele drop out (ADO) | False Allele (FA) |
| --- | --- | --- | --- | --- | --- | --- | --- | --- | --- |
| Aifu01* | 6 | 0.773 | 0.757 | 0.003 | 9.99E-02 | 3.96E-01 | -0.02 | 0.00 | 0.00 |
| CRP357* | 7 | 0.273 | 0.724 | 0.637 | 1.19E-02 | 1.66E-01 | 0.459 | 0.00 | 0.00 |
| CRP385* | 6 | 0.304 | 0.723 | 0.594 | 1.41E-03 | 6.92E-02 | 0.421 | 0.234 | 0.00 |
| CRP381* | 5 | 0.609 | 0.709 | 0.163 | 1.96E-04 | 2.98E-02 | 0.059 | 0.129 | 0.00 |
| CRP367* | 5 | 0.391 | 0.626 | 0.393 | 3.71E-05 | 1.44E-02 | 0.234 | 0.189 | 0.00 |
| CRP409* | 10 | 0.364 | 0.596 | 0.409 | 6.83E-06 | 7.19E-03 | 0.263 | 0.278 | 0.00 |
| CRP240* | 4 | 0.435 | 0.492 | 0.139 | 2.24E-06 | 4.21E-03 | 0.042 | 0.115 | 0.050 |
| CRP260 | 6 | 0.7 | 0.659 | -0.037 |  |  | -0.048 | 0.130 | 0.00 |
| Aifu05 | 5 | 0.588 | 0.694 | 0.181 |  |  | 0.046 | 0.294 | 0.00 |
| Mean | 6 | 0.493 | 0.664 | 0.276 |  |  |  |  |  |

Na—observed number of alleles, Ho—observed heterozygosity, He—expected heterozygosity, F<sub>IS</sub>— inbreeding coefficient, False alleles—null alleles, PID—probability of identity (locus), PIDsib— probability of identity for sibs (locus) is calculated with selected panel of seven loci (\* marked) according to strength of individual identification.

### Population genetic structure

The STRUCTURE analysis detected three clusters based on the mean ln P (k) and Delta K (Fig. 2 and fig. S3). We showed probabilities of population assignment at varying K (Fig. S3). At *K* - 2, a majority of individuals from WS (West Sikkim) showed a distinct cluster but no geographical assignment was observed among the individuals originated from SNP, ES (East Sikkim), and NVNP. At K-3 individuals from SNP showed a distinct isolated cluster while individuals from ES and NVNP assigned to one another cluster. However, a further increase in the K did not reveal any structuring in population. The results indicated that NVNP and ES clusters were relatively connected meta-populations as found shared ancestry in one-another populations while the other two clusters SNP and WS shared ancestry with additional genetic influxes of an unknown population which plausibly be the North Sikkim (NS) and or the historic/uncaptured gene flow from Nepal, which could not be addressed due to unavailability of samples (Fig 2a). In congruity, GENELAND also testified to the similar clustering patterns, NVNP and ES grouped into a single cluster and SNP and WS in a distinct clusters (Fig 2b). The sPCA revealed a west-east differentiation from SNP to NVNP in the allele frequencies, indicating samples of NVNP and ES in one cluster. SNP in the other and WS was assigned as an intermediate population with more affinity to SNP (Fig 2c. and Fig. S2). The DAPC also identified three major clusters, providing strong signals to support red panda populations to exist in metapopulations framework (Fig 2d). The non-Bayesian methods of population assignment in coherence to Bayesian clustering methods supported strong population genetic structure with asymmetric gene flow among the habitat patches in KL-India.

**Fig 2.**
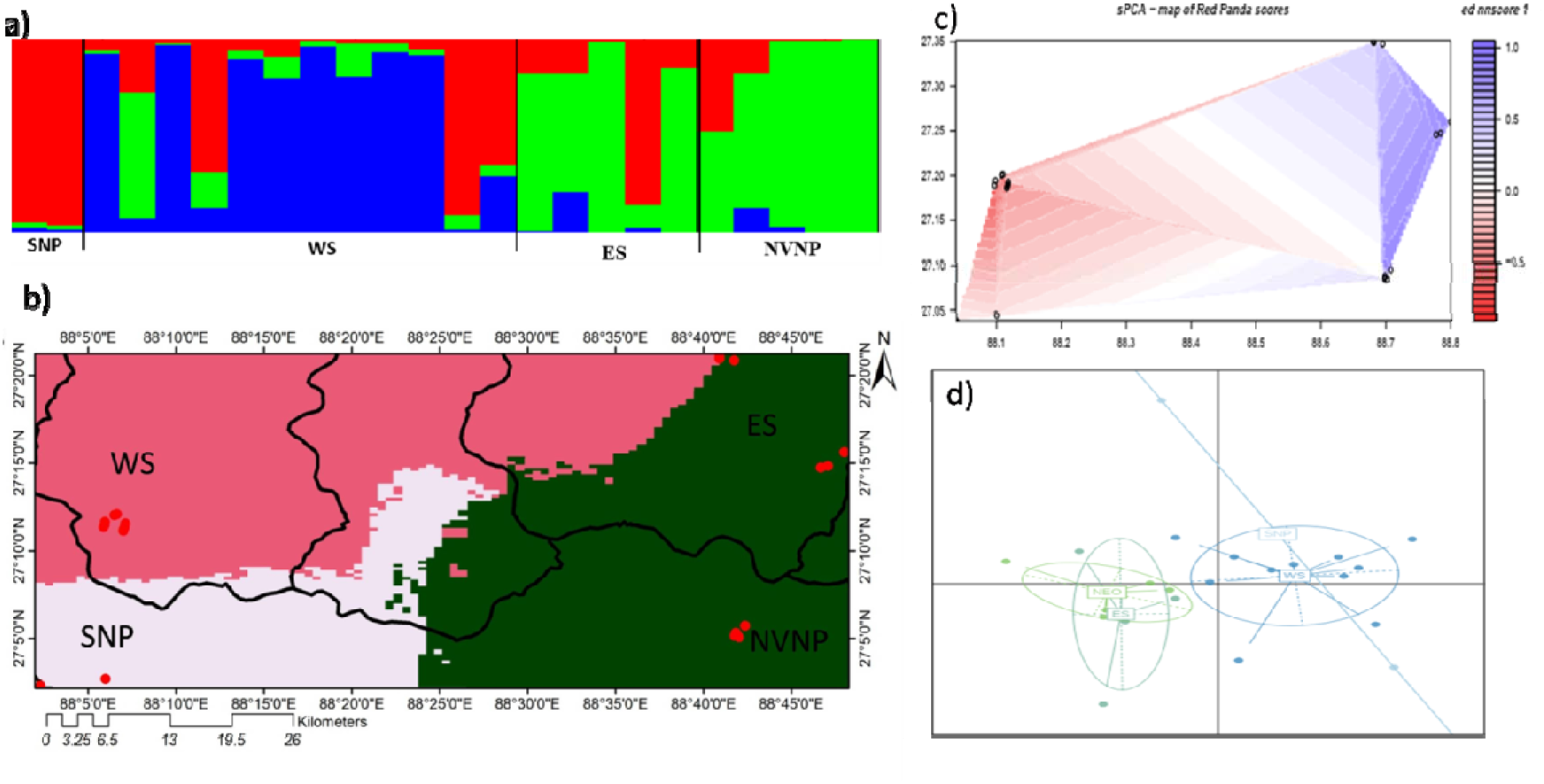
Population genetic structure of red panda population in KL-India. a) Population assignment using STRUCTURE at K3; b) Map of estimated cluster membership showing spatial distribution of the three inferred genetic clusters through GENELAND; c) Spatial PCA showing clusters in spatially distributed populations;d) Eigen values of PCA estimation showing three clusters in DAPC, each identified by individual colours and inertia eclipses.

### Gene flow and detection of migrants

In AMOVA, we found relatively lower genetic variance within population which indicated gene flow within population was higher than the between groups. Red panda populations in KL-India exhibited 58% variance within population than 12 % between populations and 30% among group with high Fst (0.321). Though Fst value was not statistically significant but observed signal was an indication of the red panda population to exist in metapopulation framework. Pairwise F_ST_ based gene flow revealed SNP and NVNP populations were highly differentiated (F_ST_ 0.352) and NVNP and ES did not qualify to be separate populations (F_ST_ −0.052) (Fig S4).

The contemporary migration assessed by the BAYESASS resulted to favour asymmetric movement of red panda from West to East in KL-India (Fig S4). We detected significant asymmetric migration of individuals from SNP to WS (5.55%) and WS to ES (5.48%) but did not obtain a rational backward gene flow. In contrary, we detected significant bidirectional migration between ES to NVNP (5.91%) and NVNP to ES (6.05%).

## Discussion

### Habitat suitability and corridor connectivity

The red panda population in KL has experienced massive habitat loss and fragmentation in the past [11,12,16, 32] which must have deteriorated connecting corridors in this landscape. For habitat suitability, predictors such as precipitation, temperature and vegetation with high weightage appeared reasonably correct as vegetation cover, and climatic factors are directly linked to the species diet, survival and reproductive necessities. A few earlier studies addressed the species distribution modelling (SDM) of Red panda in Himalaya [16, 20, 33, 34, 35]. These studies considered different variables like elevation, slope, aspect and distance to water etc. for predicting red panda habitat. These variables may facilitate in old growth forest dominated by *Betula utilis*, *Rhododendron spp.* and *Abies spp*. with dense bamboo cover in the understory and high densities of fallen logs and tree stumps at ground level. Further, the growth of bamboo understory in the temperate forests is highly influenced by the rainfall and temperature [36] and red pandas as being the canopy-dwelling species prefer temperate forests in the central and eastern Himalayas with dense bamboo undergrowth. Therefore, rainfall, temperature and vegetation cover played a significant role in predicting red panda habitat.

Earlier, Choudhury [11] estimated 1700 km^2^ potential habitat of red panda in Sikkim and 200 km^2^ in North West Bengal by undertaking exploratory field surveys based on one time efforts. Ziegler [37] estimated 684 km^2^ suitable forest for red panda in Sikkim with limited efforts confining to PWLS only. However, Ghose and Dutta [38] estimated 1200 km^2^ suitable forest for red panda in Sikkim and 250 km^2^ in North West Bengal. These authors collected primary data by undertaking field surveys through transect and trail monitoring [39] from Sikkim and North West Bengal and also recorded indirect evidences such as droppings, browsing marks, nest sites, pugmarks, skins or pelts, etc. Ghose and Dutta [38] also collected secondary data through questionnaires and public interviews in the villages around the potential red panda habitats. Our field surveys and efforts were relatively comparable to Ghose and Dutta [38] and we obtained an estimated suitable habitat 1097.26 km^2^ in Sikkim and 212.28 km^2^ in North West Bengal, which represented a relative decline the habitat suitability in last ten years when compare to Ghose and Dutta [38]. In northern region, red panda habitat was relatively fragmented and circuitscape also supported lack of horizontal connectivity in KL-India. However, the identified corridor favoured red panda movement from West to East in KL India, forming a crescent arc (Fig S4). The model demonstrated that the identified corridor in the east, south Sikkim and eastern part of north Sikkim was relatively narrow and passing through the non-PAs (Fig 1d). We propose corridor passing through the non-PAs to be monitored for the emerged developmental activities and enriched by bamboo plantation to avoid further loss of red panda habitat in KL-India.

### Genetic diversity and inbreeding

Red panda population of KL- India being a trans-boundary population, must be carrying ancestral genetic attributes shared with Nepal and Bhutan populations. The genetic assessment suggested a relatively low genetic diversity in red panda population of KL-India (H_E_= 0.66), when compared to other red panda populations i.e. H_E_= 0.719 [40] and H_E_ =0.772 [41]. The observed low genetic diversity of the Himalayan red panda might be due to historical bottlenecks [15]. The observed loss of suitable habitat by the emerged anthropogenic activities in last few decades might have disrupted the contiguous gene flow and confined red panda in isolated patches, also a contributing factor for inbreeding and loss of heterozygosity in KL-India.

### Population genetic structure and gene flow

Explicit Bayesian and non-Bayesian clustering methods to a great extent, showed similar patterns clustering red panda into at least three populations/meta-populations in KL-India. Indisputably, the NVNP and ES was found to represent one single population, However, SNP was another distinct cluster where WS was assigned as an intermediate population with more affinity to SNP. SNP and NVNP that lacks horizontal connection, were two distinct clusters (F_ST=_0.352). These populations earlier reported to exist in relatively high density (1 individual per 1.67 sq. km in SNP [20]; 32 animals in SNP and 34 in NVNP [42]). Individuals from these populations showed asymmetric gene flow and contributed migration to Sikkim population from either-side. The WS population was rationally distinct from SNP (F_ST_0.142), however ES and NVNP, to a large extent represent a single population based on the multiple evidences, e.g. a negative F_ST_-0.052 and a high migration rate from one another population (ES to NVNP-5.91% and NVNP to ES-6.05%; Fig. S4). In concurrence to this, AMOVA also supported a high variance (within group) in red panda populations. The observed scenario of contemporary gene flow patterns in KL-India suggested that corridors connecting these populations were functional in the recent past. The pragmatic asymmetric migration detected in the last two generations exhibited asymmetric migration of individuals in KL-India from West to East, SNP to WS and WS to ES. The ES and NVNP had bidirectional migration, also an indicator to support them for being a single population. The observed asymmetric gene flow among populations was due to the landscape heterogeneity and habitat suitability. The circuitscape results evident to prove that SNP and NVNP were two relatively dense populations in KL-India, where individuals might be moving from more stable, relatively high-density population to neighbouring low-density populations [43]. The high migration rate from SNP and NVNP maintained the demographic connectivity in the landscape. Both, the significant genetic differentiation in some areas and the presence of a weak population structure in other regions in KL-India may be explained by the corridor connectivity model (Fig 1d), which coincides to the results of BAYESASS and supports current flow from West to East forming a crescent arc in the KL-India.

### Functional connectivity and conservation priorities

The present study unveils the facts that landscape features have shaped the current distribution patterns of red panda in KL-India. The results demonstrated that NVNP and PWLS were connecting North West Bengal – Sikkim border. But there has been a large gap between PWL to KAS which extended up to the SRS. Further, the suitable habitat was patchy and connecting corridors were narrow between those habitat patches. Habitat on the eastern part of the study landscape was more vulnerable to destruction due to the anthropogenic activities. The FLWLS had only a small suitable area with poor connectivity to the other populations. Ziegler [37] conducted a survey in FLWLS but failed to confirm the presence of red panda. There have been recent historic records of red panda from SWLS in West Bengal [44] but intensive surveys need to be conducted to assess the current status of red panda in SWLS, which is also poorly connected to SNP.

Though, there are several PAs on the eastern part of KL-India, there is a need to extend PA network in East Sikkim, South Sikkim and eastern part of North Sikkim. Moreover, the territorial forest divisions must take adequate measures, as much of suitable habitat of red panda exists outside the PA. The present study has laid down the foundation to extend the PAs boundaries and by effectively implementing the eco-sensitive zone planning adopted by the Ministry of Environment and Forest, 2011 of Government of India [45]. We propose buffer zone may be declared around the PAs and community conservation area to protect important wildlife corridors. Since, the red panda across the range has experienced habitat loss, fragmentation and population decline due to change in the land use pattern and anthropogenic activities in past few decades. Any significant change in the climatic isotherm might result in vacating the site and or shifting the species to other sites based on varying extent of species resilience and inherent adaptive plasticity [46, 47]. Thus, red panda being ecological specialist, serves a good model to test the composite impact of landscapes, historical climate change and contemporary human activities on the possible shift in the ranges.

Further, studying wildlife using genome-wide markers (e.g. GWAS – [15, 48]; SNPs – [15, 49]) is fascinating to evaluate fine scale population genetic structure and investigating loci under natural selection facilitating populations to adapt in the changing climatic conditions [50]. However, to check the immediate effects of landscape features on the genetic variability and population contiguity, the assessment of wild populations using microsatellites is still most cost effective and widely applied way to genetic monitor of free ranging populations [30, 51]. Further, consolidating landscape connectivity through mapping corridors, validating movement through genetics and expanding natural PAs is fundamental to make red panda conservation a long-term success across the distribution range. We also seek for the possible collaboration in Nepal, Bhutan and China to aid in preparing a comprehensive monitoring plan for red panda conservation in TBL and to evaluate the ongoing conservation efforts across the political boundaries.

## Materials and methods

### Study area

KL-India, situated in the Central Himalayan biotic province with spanning between 26°21′40.49□–28°7′51.25□N and 87°30′30.67□–90°24′31.18□E. The landscape is highly rugged with mountainous terrain including the world’s 3^rd^ highest mountain peak, the Mount Kangchenjunga (8586m asl). The habitat types ranging from tropical, subtropical, warm temperate, cool temperate, subalpine, and alpine forest types [52, 53]. The study landscape is bestowed with 10 PAs *i.e.* Singalila National Park (SNP), Senchal Wildlife Sanctuary (SWLS), and Neora valley National Park (NVNP) located in North West Bengal and Barsey Rhododendron Sanctuary (BRS), Maenam Wildlife Sanctuary (MWLS), FambongLho Wildlife Sanctuary (FLWLS), Pangolakha Wildlife Sanctuary (PWLS), Kanchenjunga National Park (KNP), Shingba Rhododendron Sanctuary (SRS), and Kyongnosla Alpine Sanctuary (KAS) located in Sikkim State of India. Together these PAs, although most of them are fairly small in size, contribute approximately 3112.21 km^2^ area (Fig 1a).

### Study design and Occurrence data

Firstly, we stratified the study area based on the forest types, topography to cover all the logistically possible habitat patches in KL-India. Secondly, we adopted a landscape approach, instead of prioritizing PAs for sampling and we aimed to cover maximum reported habitats of red panda in KL-India. Thirdly, for recording the species presence, we adopted a three-pronged approach i.e. transects/ trail surveys, questionnaire surveys, and also the remote camera traps following Pradhan et al. [20], Buckland et al. [39] and Joshi et al. [54]. Further, it was not logistically feasible to cover the entire area systematically due to the rugged terrain, unpredictable weather and poor resources availability. Hence, representative sampling was carried out in KL-India that represented 109 spatially distinct locations, 12 locations from camera traps, seven direct sightings 12 presence locations from reliable interviews and 78 distinct sites where we collected 234 faecal samples of red panda. For model building, we used 56 spatially independent locations after correlation testing among the location following Kramer-Schadt [55]. Informed consent was obtained from all participants who responded to the questionnaire interviews and all methods including human participation in the study was in accordance to the relevant guidelines and regulations of Zoological Survey of India (ZSI), Kolkata. All experimental protocols and methods were approved by the Research and Academic committee (RAC) of ZSI, Kolkata. All field surveys and sampling were conducted with taking necessary permits issued by the Forest Department of West Bengal and Sikkim.

### Ecological modelling

#### Selection of environmental variables

Species are sensitive to habitats as well as the climatic isotherms [56] while, any significant change in climatic isotherm to a species having special requirements might result in local extirpation or shifting the species to other ranges [46, 47]. Therefore, it is imperative to use variables for SDM which define the likely habitat of a species based on the field observation. Considering these facts, we selected 26 variables out of which 19 variables represented climatic isotherm, four - topography and three variables represented land covers. These variables were also related to the habitat selection of red panda [16, 35,57]. The selected variables represented the present environmental conditions and habitat covariates to facilitate red panda distribution patterns in KL- India. The biotic predictors (19) were obtained from the WorldClim data base at 30 arc second scale [58]; https://www.worldclim.org/ (Table S2) and Bioclime v 1.4 dataset was used for the model building. The vegetation classification was carried out following the methodology developed by Forest Survey of India (ISFR 2017). ArcGIS 10.6 (ESRI 2018) was used to develop topographic variables from digital elevation model. For obtaining the forest canopy classes, the satellite data (Landsat 8) was downloaded from US Geological Survey (https://earthexplorer.usgs.gov/). The forest canopy was classified into four categories *viz.,* very dense, moderately dense, open forest and scrubland [59]. Moreover, considering the fact that the species is a canopy dweller [11] the Canopy height data was obtained from Oak Ridge National Laboratory (https://webmap.ornl.gov/ogc) for understanding the influence on the species habitat suitability. Since, different variables like climate, landscape, topography and anthropogenic influences used in present study were compiled from various resources present in different resolution scales. Hence, to bring uniformity in the selected data, we re-sampled all raster on the same resolution scale (1km resolution).

#### Habitat modelling and landscape connectivity

We employed maximum entropy algorithm for species distribution modelling (MaxEnt 3.4.1), using presence data only [60–62-]. Bias file was prepared in ENMeval; R package [63] following the methodology suggested by Dudik et al. [64]. We omitted highly correlated variables with a threshold of 0.8 [65, 66]. Although we selected initially 26 variables but final model was predicted using 19 variables which passed on the multicollinearity test [67] and other related variables with a threshold of 0.8 were excluded to get rid of the over fit model (Table S6) [66, 68]. For modelling, we have used 70% of the total locations as training and remaining 30% for testing the significance [60, 66, 69]. The Akaike’s information criteria (AICc) values were used to determine the best fit model with the lowest values in ENMeval. The resultant habitat was classified into suitable, with probabilities 0.584 - 1 and unsuitable below 0.584 based on the 10-percentile training presence thresholds following Radosavljevic and Anderson [70] (Table S4). The Jackknife was used to analyse the contribution of each variable provided to the MaxEnt model. We tested the model with the receiver operated characteristics area under curve (AUC) where, AUC value > 0.75 indicated high discrimination performance [71]. We also estimated the true skill statistic (TSS-0.742) that compensates for the shortcomings of kappa while keeping all of its advantages following Allouche et al. [72] We generated a raster surface of genetic divergence using Genetic landscape GIS Toolbox Ver. 10.1 [73] by generating the inverse distance weighted interpolation within the study area boundary [74]. To evaluate the operational-structural connectivity by predicted movement corridors of red panda in KL-India, we used an ensemble approach by combining both the genetic divergence and environmental conductance (SDM output) in the final circuit model following Mateo-Sánchez et. al.[75] and Roffler et.al [76] in Circuitscape 4.0 [77].

### Population genetic analyses

#### DNA extraction, PCR amplification and microsatellite genotyping

We collected 234 red panda feces, i.e. 87 feces from North West Bengal and 147 from Sikkim in KL- India (Fig.1a). All samples were stored in 70% ethanol and DNA was extracted using QIAamp Fast DNA Stool Mini Kit (QIAGEN Germany) following manufacturer’s instructions. Nine polymorphic STRs, *i.e.* Aifu01, Aifu05 from Liang [78] and CRP357, CRP385, CRP381, CRP367, CRP409, CRP240 and CRP260 from Yang [79] were amplified into two multiplex PCRs. Forward primer of all nine microsatellite loci was fluorescently labelled at 5’ with one of the four dyes, FAM, VIC, NED and PET (Table S5). The PCRs were carried-out in 10μl reaction volume following QIAGEN Multiplex PCR Kit (Qiagen, Germany). The thermal cycle profile was: initial denaturation at 95°C for 15 minutes, followed by 40 cycles of PCR and a final step of 72° C for 30 minutes. The annealing temperature (Ta) for multiplex 1 (CRP381, CRP367, CRP409, CRP240 and CRP260) was 55 °C and multiplex 2 (Aifu01, Aifu05, CRP357, CRP385) was 57 °C (Table S5). The PCR products were resolved on an ABI 3730 Genetic analyzer (Applied Biosystems, Foster City, CA, USA) and allele scoring was done using Gene Mapper 4.1 (Applied Biosystems, Foster City, CA, USA).

#### Genotyping error and individual identification

We genotyped each sample four times to minimize genotyping errors and a heterozygote was ascertained only if there were different alleles in at least three independent attempts. In addition, we also followed an independent allele scoring method where two different researchers perform allele scoring individually and only the consensus genotypes were used for further analysis. The genotyping errors arising due to null allele and the presence of stutters, scoring errors were assessed using MICRO CHECKER 2.2.2 [80]. Maximum likelihood allele dropout (ADO) and false allele (FA) error rates were quantified using PEDANT version 1.0 involving 10,000 search steps for enumeration of per allele error rates [81]. To avoid ambiguity in ascertaining unique genotypes, we limited the number of loci used based on their high success rate (>90%), presence of no or minimum genotyping errors and exhibiting an informative *P_ID_* value (probability of obtaining identical genotypes between two samples by chance). The locus wise and cumulative probability of identity for unrelated individuals (*P_ID_*) and siblings (*P_ID_ sibs*) were calculated following identity analysis module in GenAlEx version 6 [82].

### Genetic diversity and inbreeding

The genetic diversity estimates were accounted by calculating the number of allele (Na), observed (Ho) and expected (He) heterozygosity using GENALEX 6 [82]. For the Hardy-Weinberg equilibrium (HWE) test, we followed the probability test approach using the program GENEPOP version 4.2.1 [83]. Wright’s inbreeding coefficient (*F_IS_*) was estimated following Weir & Cockerham [84] using GENEPOP [83]. Linkage disequilibrium (LD) was tested using GENEPOP [83] to determine the extent of distortion from independent segregation of loci following 10,000 dememorizations, 500 batches and 10,000 iterations per batch after Bonferroni correction [85].

### Population genetic structure

We attempted three different clustering methods to capture the most possible population genetic structure of red panda in KL-India *i.e.* Fst based Analysis of Molecular Variance (AMOVA), explicit Bayesian and non-Bayesian clustering algorithms. We used Arlequin 3.5.2.1 [86] to estimate the proportions of the total genetic variation, arose from, within and between populations using AMOVA. Among different Bayesian clustering methods, individuals were assigned exclusively on the basis of their multi-locus genotypes (e.g. STRUCTURE), and also using the both, multi-locus genotypes and geo-referenced information (e.g. GENELAND). Non-Bayesian multivariate ordination analyses, i.e. discriminant analysis of principle components (DAPC) and spatial principle component analysis (sPCA) were also used, to compare population assignment with the Bayesian clustering outputs [87] since they were not based on any model assumptions.

In Bayesian analysis, STRUCTURE 2.3.4 software [88] was used to determine the number of genetic clusters (K) following 20 iterations (20,000 burn-in; 200,000 Markov chain Monte Carlo replicates in each run) with NOPRIOR with admixed and correlated allele frequencies. We considered there were K populations (1 to 10), with repeating each analysis for 10 times at each K value. The most probable cluster was calculated via estimating the distribution of Delta K [89] using STRUCTURE HARVESTER v.0.68 [90]. GENELAND v4.0.3 [91] was run through an extension of R v.3.0.1 with the correlated allele frequency and spatial uncertainty model. We allowed K to vary between 1 to 10 following 20 independent runs, each with 100000 iterations, and a thinning of 1000. The inferred spatial clusters were georeferenced in ArcGIS 10.6. Two non-Bayesian clustering methods *i.e.*, DAPC and sPCA, were run to assign the possible clusters in Adegenet v1.3.4 package of R [92].

### Gene flow and migration rate

We assessed Fst based gene flow in last 150-200 years [93] among the different sub-populations of red panda in KL-India using Arlequin v 3.5.2 [86]. The contemporary and asymmetric migration rate in last two generations were estimated using BayesAss 1.3 [94]. We used 9×10^6^ iterations, with a burn-in of 10^6^ iterations, 1000 number of permutations and a sampling frequency of 2000 to ensure that the model’s starting parameters were sufficiently randomized. We also estimated the first-generation migrants between all pairs of subpopulations of red panda in KL-India using GENECLASS 2.0 [95], an approach described by Paetkau [96] for likelihood computation (Lhome/Lmax), with 1000 simulations at an assignment threshold (alpha) of 0.01 and 0.05.

## ACKNOWLEDGEMENTS

Authors thank Principal Chief Wildlife Warden, Forest Departments of red panda range States for granting the necessary permission to carry out the study. The study was supported by the DST - INSPIRE FACULTY SCHEME (Grant No. DST/INSPIRE/04/2016/002246) awarded to Dr Mukesh Thakur and funded received by the NMHS-Large Grant project of the Ministry of Environment, Forest and Climate Change, New Delhi (Grant No. NMHS/2017-18/LG09/02).

