## Supplementary files for "Fine-scale landscape genetics unveiling contemporary asymmetric movement of red panda (*Ailurus fulgens*) in Kangchenjunga landscape, India"

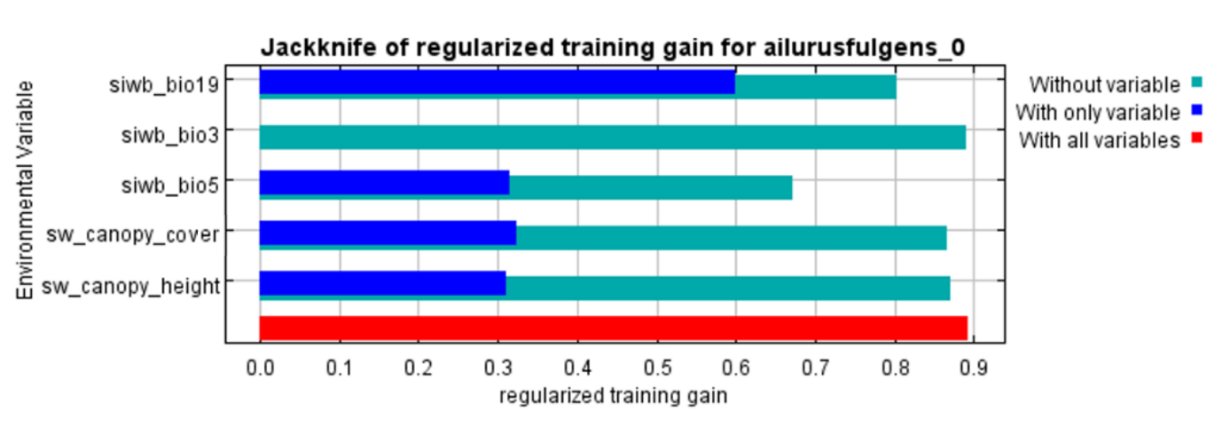
Supplementary Figures and Tables

Fig S1: Jackknife test for all five variables. The environmental variable with highest gain when used in isolation is siwb_bio19, therefore it has the most useful information. The environmental variable that decreases the gain the most when it is omitted is siwb_bio5, which therefore appears to have the most information that isn't present in the other variables.

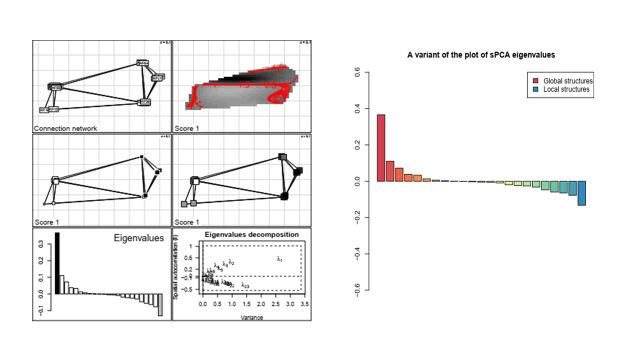
Fig. S2: Spatial PCA result showing clusters in spatially distributed populations

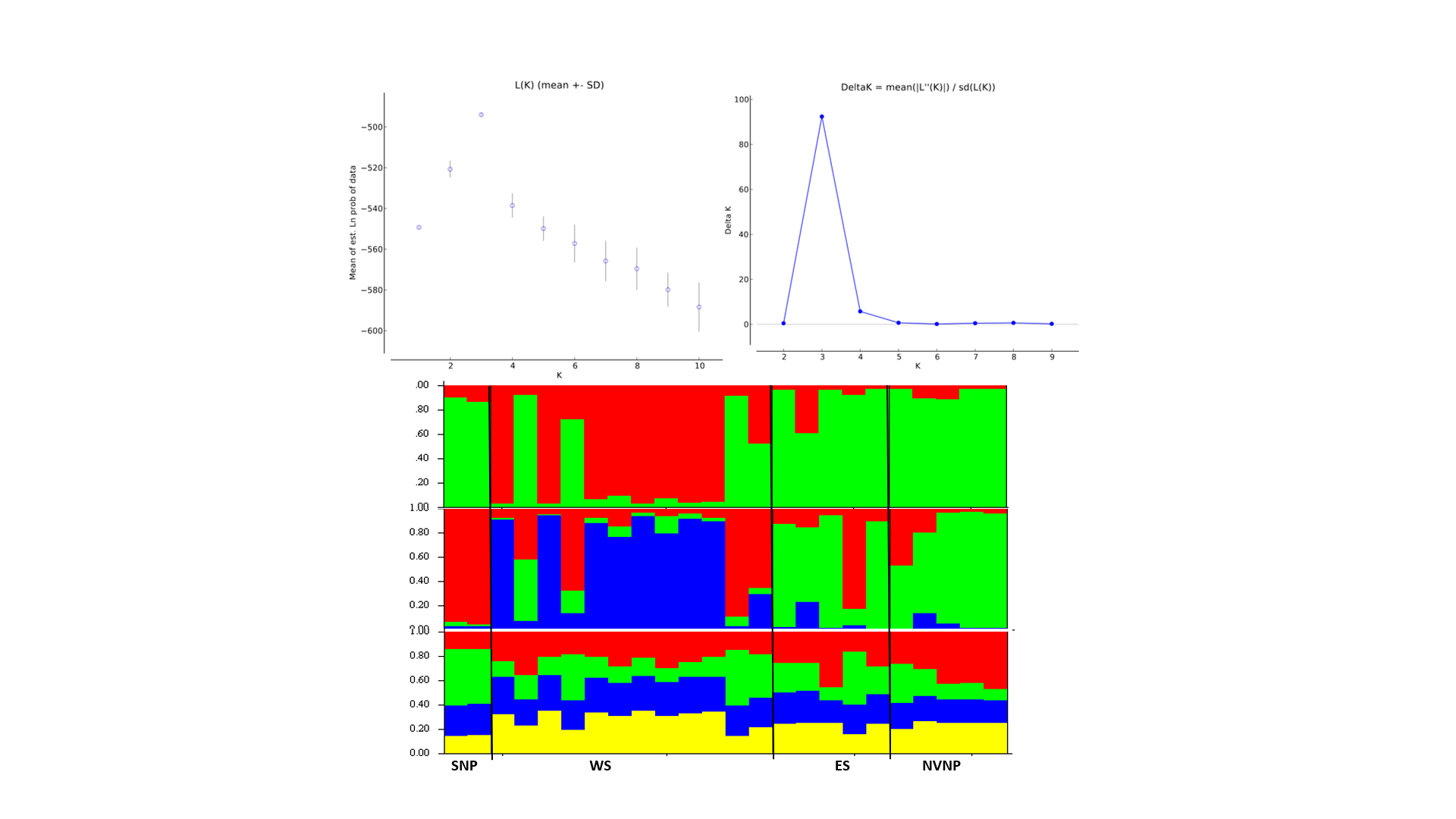

K4

K3

K2

c)

a)

b)

Fig. S3: Population assignment probabilities of red panda populations based on STRUCTURE in KL-India. a) lnP(K) (posterior probability of the data), b) Magnitude of delta K (rate of change in the log probability) and c) Summary bar plots of STRUCTURE run from K2 to K 4

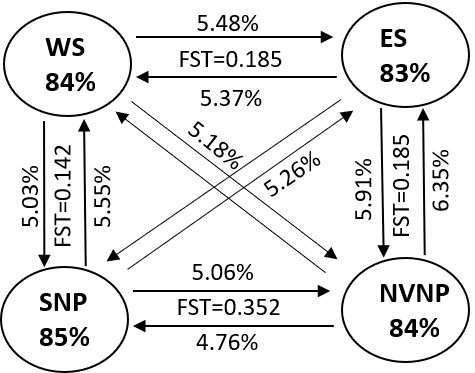

Fig. S4: Schametic diagram of Contemporary migration rate (> 5% as significant value and FST among red panda populations SNP- Singalila National Park. WS: West Sikkim (Composite landscape comprising of Barsey Rhododendron Wildlife Sanctuary and Kanchenjunga National Park, ES: East Sikkim (Composite landscpe of Pangolakha Wildlife Sanctuary and Kyongnosola Alpine sanctuary and NVNP: Neora Valley National Park)

Table S1: List of all predictor variables

| Variable | Code |
| --- | --- |
| Annual Mean Temperature | siwb_bio1 |
| Mean Diurnal Range (Mean of monthly (max temp - min temp) | siwb_bio2 |
| Isothermality (BIO2/BIO7) | siwb_bio3 |
| Temperature Seasonality (Standard Deviation) | siwb_bio4 |
| Max Temperature of Warmest Month | siwb_bio5 |
| Min Temperature of Coldest Month | siwb_bio6 |
| Temperature Annual Range (BIO5-BIO6) | siwb_bio7 |
| Mean Temperature of Wettest Quarter | siwb_bio8 |
| Mean Temperature of Driest Quarter | siwb_bio9 |
| Mean Temperature of Warmest Quarter | siwb_bio10 |
| Mean Temperature of Coldest Quarter | siwb_bio11 |
| Annual Precipitation | siwb_bio12 |
| Precipitation of Wettest Month | siwb_bio13 |
| Precipitation of Driest Month | siwb_bio14 |
| Precipitation Seasonality (Coefficient of Variation) | siwb_bio15 |
| Precipitation of Wettest Quarter | siwb_bio16 |
| Precipitation of Driest Quarter | siwb_bio17 |
| Precipitation of Warmest Quarter | siwb_bio18 |
| Precipitation of Coldest Quarter | siwb_bio19 |
| Canopy Cover | sw_canopy_cover |
| Canopy Height | sw_canopy_height |
| Normalized Difference Vegetation Index | siwb_ndvi |
| Roughness | siwb_rough |
| Slope | siwb_slope |
| Hillshade | siwb_hills |
| Aspect | siwb_aspec |

Table S2: Correlation matrix between least correlated variables

| ***Column1*** | ***siwb_bio3*** | ***siwb_bio5*** | ***siwb_bio19*** | ***siwb_lulc_*** | ***sw_canopy_*** |
| --- | --- | --- | --- | --- | --- |
| siwb_bio3 | 1.00 |  |  |  |  |
| siwb_bio5 | -0.58 | 1.00 |  |  |  |
| siwb_bio19 | 0.25 | -0.17 | 1.00 |  |  |
| sw_canopy_cover | -0.05 | 0.03 | 0.07 | 1.00 |  |
| sw_canopy_height | -0.13 | -0.15 | 0.01 | -0.36 | 1.00 |

Table S3: Best fit model setting selected on the basis of lowest AICc value inENMEval.

| S.N. |  | **Settings** | **features** | **rm** | **train. AUC** | **avg.test. AUC** | **var.test. AUC** |  | **avg.diff. AUC** | **var.diff. AUC** | **avg.test. orMTP** | **var.test. orMTP** | **avg.test. or10pct** | **var.test. or10pct** | **AICc** | **delta. AICc** | **w.AIC** | **param eters** |
| --- | --- | --- | --- | --- | --- | --- | --- | --- | --- | --- | --- | --- | --- | --- | --- | --- | --- | --- |
| 1 |  | L_0.5 | L | 0.5 | 0.93 | 0.88 | 0.44 |  | 0.06 | 0.35 | 0.03 | 0.03 | 0.26 | 0.2 | 526.51 | 27.81 | 0 | 14 |
| 2 |  | LQ_0.5 | LQ | 0.5 | 0.93 | 0.88 | 0.47 |  | 0.07 | 0.4 | 0.06 | 0.06 | 0.23 | 0.18 | 558.59 | 59.88 | 0 | 18 |
| 3 |  | H_0.5 | H | 0.5 | 0.96 | 0.88 | 0.42 |  | 0.09 | 0.38 | 0.16 | 0.14 | 0.58 | 0.25 | NA | NA | NA | 55 |
| 4 |  | LQH_0.5 | LQH | 0.5 | 0.96 | 0.88 | 0.45 |  | 0.09 | 0.41 | 0.13 | 0.12 | 0.52 | 0.26 | NA | NA | NA | 57 |
| 5 |  | LQHP_0.5 | LQHP | 0.5 | 0.96 | 0.88 | 0.4 |  | 0.09 | 0.36 | 0.19 | 0.16 | 0.58 | 0.25 | NA | NA | NA | 57 |
| 6 |  | LQHPT_0.5 | LQHPT | 0.5 | 0.96 | 0.88 | 0.4 |  | 0.09 | 0.36 | 0.19 | 0.16 | 0.58 | 0.25 | NA | NA | NA | 57 |
| 7 |  | L_1 | L | 1 | 0.92 | 0.89 | 0.47 |  | 0.06 | 0.38 | 0.03 | 0.03 | 0.26 | 0.2 | 517.15 | 18.45 | 0 | 12 |
| 8 |  | LQ_1 | LQ | 1 | 0.93 | 0.89 | 0.46 |  | 0.06 | 0.38 | 0.03 | 0.03 | 0.23 | 0.18 | 513.53 | 14.82 | 0 | 12 |
| 9 |  | H_1 | H | 1 | 0.95 | 0.89 | 0.35 |  | 0.07 | 0.3 | 0.06 | 0.06 | 0.29 | 0.21 | 2247.81 | 1749.1 | 0 | 29 |
| 10 |  | LQH_1 | LQH | 1 | 0.94 | 0.88 | 0.46 |  | 0.07 | 0.4 | 0.13 | 0.12 | 0.26 | 0.2 | NA | NA | NA | 33 |
| 11 |  | LQHP_1 | LQHP | 1 | 0.95 | 0.88 | 0.4 |  | 0.08 | 0.35 | 0.13 | 0.12 | 0.26 | 0.2 | NA | NA | NA | 32 |
| 12 |  | LQHPT_1 | LQHPT | 1 | 0.95 | 0.88 | 0.4 |  | 0.08 | 0.35 | 0.13 | 0.12 | 0.26 | 0.2 | NA | NA | NA | 32 |
| 13 |  | L_1.5 | L | 1.5 | 0.92 | 0.89 | 0.48 |  | 0.06 | 0.39 | 0.03 | 0.03 | 0.23 | 0.18 | 509.68 | 10.97 | 0 | 10 |
| 14 |  | LQ_1.5 | LQ | 1.5 | 0.92 | 0.89 | 0.51 |  | 0.06 | 0.42 | 0.03 | 0.03 | 0.19 | 0.16 | 512.43 | 13.73 | 0 | 11 |
| 15 |  | H_1.5 | H | 1.5 | 0.94 | 0.9 | 0.32 |  | 0.06 | 0.26 | 0.06 | 0.06 | 0.26 | 0.2 | 533.72 | 35.01 | 0 | 16 |
| 16 |  | LQH_1.5 | LQH | 1.5 | 0.93 | 0.89 | 0.45 |  | 0.07 | 0.38 | 0.13 | 0.12 | 0.19 | 0.16 | 558.89 | 60.19 | 0 | 18 |
| 17 |  | LQHP_1.5 | LQHP | 1.5 | 0.94 | 0.88 | 0.45 |  | 0.07 | 0.38 | 0.13 | 0.12 | 0.23 | 0.18 | 573.97 | 75.26 | 0 | 19 |
| 18 |  | LQHPT_1.5 | LQHPT | 1.5 | 0.94 | 0.88 | 0.45 |  | 0.07 | 0.38 | 0.13 | 0.12 | 0.23 | 0.18 | 573.97 | 75.26 | 0 | 19 |
| 19 |  | L_2 | L | 2 | 0.92 | 0.89 | 0.52 |  | 0.05 | 0.42 | 0.03 | 0.03 | 0.23 | 0.18 | 508.41 | 9.7 | 0 | 9 |
| 20 |  | LQ_2 | LQ | 2 | 0.92 | 0.89 | 0.54 |  | 0.06 | 0.44 | 0.03 | 0.03 | 0.19 | 0.16 | 503.35 | 4.65 | 0.02 | 8 |
| 21 |  | H_2 | H | 2 | 0.94 | 0.9 | 0.3 |  | 0.06 | 0.24 | 0.06 | 0.06 | 0.19 | 0.16 | 518.25 | 19.55 | 0 | 13 |
| 22 |  | LQH_2 | LQH | 2 | 0.93 | 0.89 | 0.46 |  | 0.06 | 0.37 | 0.06 | 0.06 | 0.23 | 0.18 | 504.66 | 5.96 | 0.01 | 10 |
| 23 |  | LQHP_2 | LQHP | 2 | 0.93 | 0.88 | 0.54 |  | 0.07 | 0.45 | 0.06 | 0.06 | 0.19 | 0.16 | 537.66 | 38.95 | 0 | 15 |
| 24 |  | LQHPT_2 | LQHPT | 2 | 0.93 | 0.88 | 0.54 |  | 0.07 | 0.45 | 0.06 | 0.06 | 0.19 | 0.16 | 537.66 | 38.95 | 0 | 15 |
| 25 |  | L_2.5 | L | 2.5 | 0.91 | 0.89 | 0.56 |  | 0.06 | 0.45 | 0.03 | 0.03 | 0.23 | 0.18 | 504.07 | 5.37 | 0.02 | 7 |
| 26 |  | LQ_2.5 | LQ | 2.5 | 0.92 | 0.9 | 0.52 |  | 0.05 | 0.43 | 0.03 | 0.03 | 0.19 | 0.16 | 501.07 | 2.36 | 0.07 | 7 |
| 27 |  | H_2.5 | H | 2.5 | 0.93 | 0.9 | 0.31 |  | 0.06 | 0.23 | 0.1 | 0.09 | 0.19 | 0.16 | 514.3 | 15.6 | 0 | 11 |
| 28 |  | LQH_2.5 | LQH | 2.5 | 0.92 | 0.89 | 0.51 |  | 0.06 | 0.41 | 0.06 | 0.06 | 0.19 | 0.16 | 505.28 | 6.57 | 0.01 | 9 |
| 29 |  | LQHP_2.5 | LQHP | 2.5 | 0.93 | 0.88 | 0.64 |  | 0.07 | 0.54 | 0.06 | 0.06 | 0.19 | 0.16 | 517.28 | 18.57 | 0 | 11 |
| 30 |  | LQHPT_2.5 | LQHPT | 2.5 | 0.93 | 0.88 | 0.64 |  | 0.07 | 0.54 | 0.06 | 0.06 | 0.19 | 0.16 | 517.28 | 18.57 | 0 | 11 |
| 31 |  | L_3 | L | 3 | 0.91 | 0.89 | 0.56 |  | 0.05 | 0.44 | 0.03 | 0.03 | 0.23 | 0.18 | 502.01 | 3.31 | 0.05 | 6 |
| 32 |  | LQ_3 | LQ | 3 | 0.92 | 0.9 | 0.51 |  | 0.05 | 0.41 | 0.03 | 0.03 | 0.19 | 0.16 | 498.7 | 0 | 0.24 | 6 |
| 33 |  | H_3 | H | 3 | 0.93 | 0.89 | 0.31 |  | 0.05 | 0.23 | 0.03 | 0.03 | 0.19 | 0.16 | 502.98 | 4.27 | 0.03 | 7 |
| 34 |  | LQH_3 | LQH | 3 | 0.92 | 0.89 | 0.51 |  | 0.05 | 0.42 | 0.03 | 0.03 | 0.19 | 0.16 | 501.24 | 2.53 | 0.07 | 7 |
| 35 |  | LQHP_3 | LQHP | 3 | 0.92 | 0.88 | 0.69 |  | 0.07 | 0.59 | 0.06 | 0.06 | 0.19 | 0.16 | 507.82 | 9.11 | 0 | 8 |
| 36 |  | LQHPT_3 | LQHPT | 3 | 0.92 | 0.88 | 0.69 |  | 0.07 | 0.59 | 0.06 | 0.06 | 0.19 | 0.16 | 507.82 | 9.11 | 0 | 8 |
| 37 |  | L_3.5 | L | 3.5 | 0.91 | 0.89 | 0.56 |  | 0.05 | 0.44 | 0.03 | 0.03 | 0.19 | 0.16 | 502.79 | 4.09 | 0.03 | 6 |
| 38 |  | LQ_3.5 | LQ | 3.5 | 0.92 | 0.9 | 0.51 |  | 0.05 | 0.41 | 0.03 | 0.03 | 0.19 | 0.16 | 499.82 | 1.12 | 0.14 | 6 |
| 39 |  | H_3.5 | H | 3.5 | 0.92 | 0.89 | 0.34 |  | 0.05 | 0.24 | 0.03 | 0.03 | 0.26 | 0.2 | 508.37 | 9.67 | 0 | 7 |
| 40 |  | LQH_3.5 | LQH | 3.5 | 0.92 | 0.89 | 0.51 |  | 0.05 | 0.41 | 0.03 | 0.03 | 0.19 | 0.16 | 499.82 | 1.12 | 0.14 | 6 |
| 41 |  | LQHP_3.5 | LQHP | 3.5 | 0.92 | 0.88 | 0.73 |  | 0.06 | 0.62 | 0.06 | 0.06 | 0.23 | 0.18 | 511.33 | 12.63 | 0 | 8 |
| 42 |  | LQHPT_3.5 | LQHPT | 3.5 | 0.92 | 0.88 | 0.73 |  | 0.06 | 0.62 | 0.06 | 0.06 | 0.23 | 0.18 | 511.33 | 12.63 | 0 | 8 |
| 43 |  | L_4 | L | 4 | 0.91 | 0.89 | 0.56 |  | 0.06 | 0.45 | 0.03 | 0.03 | 0.19 | 0.16 | 503.65 | 4.95 | 0.02 | 6 |
| 44 |  | LQ_4 | LQ | 4 | 0.92 | 0.89 | 0.51 |  | 0.05 | 0.42 | 0.03 | 0.03 | 0.16 | 0.14 | 501.05 | 2.35 | 0.07 | 6 |
| 45 |  | H_4 | H | 4 | 0.91 | 0.89 | 0.36 |  | 0.05 | 0.26 | 0.03 | 0.03 | 0.23 | 0.18 | 513.8 | 15.09 | 0 | 7 |
| 46 |  | LQH_4 | LQH | 4 | 0.92 | 0.89 | 0.51 |  | 0.05 | 0.42 | 0.03 | 0.03 | 0.16 | 0.14 | 501.05 | 2.35 | 0.07 | 6 |
| 47 |  | LQHP_4 | LQHP | 4 | 0.92 | 0.88 | 0.75 |  | 0.06 | 0.64 | 0.06 | 0.06 | 0.23 | 0.18 | 507.53 | 8.82 | 0 | 6 |
| 48 |  | LQHPT_4 | LQHPT | 4 | 0.92 | 0.88 | 0.75 |  | 0.06 | 0.64 | 0.06 | 0.06 | 0.23 | 0.18 | 507.53 | 8.82 | 0 | 6 |

Table S4: Threshold selection for MaxEnt model at 10 percentile training presence

| **Cumulative threshold** | **Cloglog threshold** | **Description** | **Fractional predicted area** | **Training omission rate** | **Test omission rate** | **p-value** |
| --- | --- | --- | --- | --- | --- | --- |
| 1.000 | 0.060 | Fixed cumulative value 1 | 0.687 | 0.000 | 0.000 | 6.868E^-1^ |
| 5.000 | 0.116 | Fixed cumulative value 5 | 0.540 | 0.000 | 0.000 | 5.403E^-1^ |
| 10.000 | 0.153 | Fixed cumulative value 10 | 0.410 | 0.000 | 0.000 | 4.097E^-1^ |
| 18.785 | 0.256 | Minimum training presence | 0.260 | 0.000 | 0.000 | 2.595E^-1^ |
| 36.553 | 0.568 | 10 percentile training presence | 0.125 | 0.100 | 0.100 | 1.255E^-1^ |
| 36.576 | 0.568 | Equal training sensitivity and specificity | 0.125 | 0.133 | 0.000 | 1.255E^-1^ |
| 35.198 | 0.544 | Maximum training sensitivity plus specificity | 0.132 | 0.067 | 0.000 | 1.317E^-1^ |
| 40.320 | 0.624 | Equal training sensitivity and specificity | 0.110 | 0.233 | 0.000 | 1.101E^-1^ |
| 40.320 | 0.624 | Maximum training sensitivity plus specificity | 0.110 | 0.233 | 0.000 | 1.101E^-1^ |
| 8.118 | 0.138 | Balance training omission, predicted area and threshold value | 0.545 | 0.000 | 0.000 | 4.544E^-1^ |
| 11.795 | 0.168 | Equate entropy of threshold and original distribution | 0.372 | 0.000 | 0.000 | 3.716E^-1^ |

Table S5: Characteristics of microsatellite loci used in the present study

| **S. No.** | **Locus** | **Dye** | **Size Range** | **Repeat Motif** | **Ta (°C)** | **Multiplex** | **Forward primer (5'--3')** | **Reverse primer (5'--3')** | **Reference** |
| --- | --- | --- | --- | --- | --- | --- | --- | --- | --- |
| 1 | CRP260 | VIC | 200-250 | Tetra | 61 | MP1 | GGGGCCTTGTCTAATTCTGT | TCACTCCTGGTGCTGGTCT | Yang et al. 2019 |
| 2 | Aifu01 | FAM | 126-178 | Tetra | 61 |  | CCTGCATCAGACTCAGCA | GGTATCAGACGTGGGAACTA | Liang et al. 2007 |
| 3 | Aifu05 | FAM | 328-348 | Tetra | 61 |  | GAATAATGAGCTTGCCTTCC | TTGACATTGGCTATGTGAACA | Liang et al. 2007 |
| 4 | CRP357 | VIC | 360-410 | Tri | 53 | MP2 | TCCAAAATAATGGTAAAGCC | CAACTCAACTACATCGCCTC | Yang et al. 2019 |
| 5 | CRP381 | PET | 240-290 | Tetra | 53 |  | AATGTCAAGGAAGAACCCAA | TTCACTGCTCACCGTTTCTA | Yang et al. 2019 |
| 6 | CRP367 | VIC | 240-290 | Tetra | 53 |  | TCCATGTAAGCCTCCAAACT | AGAACCAAATGTCTCCACGT | Yang et al. 2019 |
| 7 | CRP385 | FAM | 370-420 | Tetra | 53 |  | CATCCCAGGAGACCAAAG | ATAAACTGACAAGAAGTCCTCC | Yang et al. 2019 |
| 8 | CRP240 | VIC | 160-210 | Tetra | 55 |  | TGATTCAAGGTTCCCTATGT | AAGAAAGTGGTTAGTTTAAATGTT | Yang et al. 2019 |
| 9 | CRP409 | NED | 270-320 | Tri | 55 |  | CCACCATCTGTTAGGGAGTAG | CCTTGATTTGTTGGAGCATT | Yang et al. 2019 |

Table S6: Correlation matrix of *Topographic variable used in the present study.*

| ***Column1*** | ***siwb_bio1*** | ***siwb_bio2*** | ***siwb_bio3*** | ***siwb_bio4*** | ***siwb_bio5*** | ***siwb_bio6*** | ***siwb_bio7*** | ***siwb_bio8*** | ***siwb_bio9*** | ***siwb_bio10*** | ***siwb_bio11*** | ***siwb_bio12*** | ***siwb_bio13*** | ***siwb_bio14*** | ***siwb_bio15*** | ***siwb_bio16*** | ***siwb_bio17*** | ***siwb_bio18*** | ***siwb_bio19*** | ***siwb_lulc_*** | ***sw_canopy_*** | ***siwb_ndvi*** | ***siwb_rough*** | ***siwb_slope*** | ***siwb_hills*** | ***siwb_aspec*** |
| --- | --- | --- | --- | --- | --- | --- | --- | --- | --- | --- | --- | --- | --- | --- | --- | --- | --- | --- | --- | --- | --- | --- | --- | --- | --- | --- |
| siwb_bio1 | 1.00 |  |  |  |  |  |  |  |  |  |  |  |  |  |  |  |  |  |  |  |  |  |  |  |  |  |
| siwb_bio2 | -0.87 | 1.00 |  |  |  |  |  |  |  |  |  |  |  |  |  |  |  |  |  |  |  |  |  |  |  |  |
| siwb_bio3 | -0.82 | 0.86 | 1.00 |  |  |  |  |  |  |  |  |  |  |  |  |  |  |  |  |  |  |  |  |  |  |  |
| siwb_bio4 | -0.78 | 0.96 | 0.70 | 1.00 |  |  |  |  |  |  |  |  |  |  |  |  |  |  |  |  |  |  |  |  |  |  |
| siwb_bio5 | 0.86 | -0.50 | -0.58 | -0.38 | 1.00 |  |  |  |  |  |  |  |  |  |  |  |  |  |  |  |  |  |  |  |  |  |
| siwb_bio6 | 0.97 | -0.96 | -0.83 | -0.91 | 0.71 | 1.00 |  |  |  |  |  |  |  |  |  |  |  |  |  |  |  |  |  |  |  |  |
| siwb_bio7 | -0.85 | 0.99 | 0.80 | 0.99 | -0.46 | -0.95 | 1.00 |  |  |  |  |  |  |  |  |  |  |  |  |  |  |  |  |  |  |  |
| siwb_bio8 | 0.97 | -0.74 | -0.77 | -0.63 | 0.95 | 0.89 | -0.71 | 1.00 |  |  |  |  |  |  |  |  |  |  |  |  |  |  |  |  |  |  |
| siwb_bio9 | 0.99 | -0.92 | -0.82 | -0.85 | 0.80 | 0.99 | -0.90 | 0.94 | 1.00 |  |  |  |  |  |  |  |  |  |  |  |  |  |  |  |  |  |
| siwb_bio10 | 0.97 | -0.74 | -0.77 | -0.63 | 0.95 | 0.89 | -0.71 | 1.00 | 0.94 | 1.00 |  |  |  |  |  |  |  |  |  |  |  |  |  |  |  |  |
| siwb_bio11 | 0.99 | -0.91 | -0.82 | -0.85 | 0.80 | 0.99 | -0.90 | 0.94 | 1.00 | 0.94 | 1.00 |  |  |  |  |  |  |  |  |  |  |  |  |  |  |  |
| siwb_bio12 | 0.96 | -0.95 | -0.88 | -0.88 | 0.71 | 0.98 | -0.93 | 0.89 | 0.98 | 0.89 | 0.98 | 1.00 |  |  |  |  |  |  |  |  |  |  |  |  |  |  |
| siwb_bio13 | 0.95 | -0.95 | -0.90 | -0.86 | 0.69 | 0.97 | -0.93 | 0.88 | 0.97 | 0.88 | 0.96 | 1.00 | 1.00 |  |  |  |  |  |  |  |  |  |  |  |  |  |
| siwb_bio14 | 0.09 | -0.37 | 0.06 | -0.57 | -0.24 | 0.28 | -0.46 | -0.08 | 0.18 | -0.08 | 0.19 | 0.16 | 0.15 | 1.00 |  |  |  |  |  |  |  |  |  |  |  |  |
| siwb_bio15 | 0.88 | -0.94 | -0.86 | -0.87 | 0.59 | 0.93 | -0.92 | 0.79 | 0.91 | 0.79 | 0.91 | 0.96 | 0.98 | 0.19 | 1.00 |  |  |  |  |  |  |  |  |  |  |  |
| siwb_bio16 | 0.95 | -0.95 | -0.89 | -0.87 | 0.69 | 0.97 | -0.93 | 0.88 | 0.97 | 0.88 | 0.96 | 1.00 | 1.00 | 0.14 | 0.98 | 1.00 |  |  |  |  |  |  |  |  |  |  |
| siwb_bio17 | 0.47 | -0.67 | -0.26 | -0.84 | 0.11 | 0.63 | -0.75 | 0.31 | 0.56 | 0.30 | 0.56 | 0.52 | 0.49 | 0.89 | 0.49 | 0.49 | 1.00 |  |  |  |  |  |  |  |  |  |
| siwb_bio18 | 0.95 | -0.95 | -0.89 | -0.87 | 0.69 | 0.98 | -0.93 | 0.88 | 0.97 | 0.88 | 0.97 | 1.00 | 1.00 | 0.16 | 0.97 | 1.00 | 0.50 | 1.00 |  |  |  |  |  |  |  |  |
| siwb_bio19 | 0.04 | -0.18 | 0.25 | -0.42 | -0.17 | 0.17 | -0.29 | -0.09 | 0.11 | -0.09 | 0.11 | 0.03 | -0.03 | 0.83 | -0.06 | -0.03 | 0.82 | -0.01 | 1.00 |  |  |  |  |  |  |  |
| siwb_lulc_ | 0.00 | 0.01 | -0.05 | 0.03 | 0.03 | -0.01 | 0.02 | 0.01 | 0.00 | 0.01 | -0.01 | -0.03 | -0.04 | 0.00 | -0.07 | -0.04 | 0.00 | -0.06 | 0.07 | 1.00 |  |  |  |  |  |  |
| sw_canopy_ | 0.00 | -0.15 | -0.13 | -0.16 | -0.15 | 0.07 | -0.16 | -0.06 | 0.03 | -0.06 | 0.03 | 0.09 | 0.11 | 0.17 | 0.14 | 0.11 | 0.14 | 0.10 | 0.01 | -0.36 | 1.00 |  |  |  |  |  |
| siwb_ndvi | 0.22 | -0.18 | -0.33 | -0.07 | 0.21 | 0.18 | -0.14 | 0.24 | 0.20 | 0.24 | 0.19 | 0.23 | 0.25 | -0.26 | 0.21 | 0.25 | -0.16 | 0.23 | -0.30 | 0.21 | -0.06 | 1.00 |  |  |  |  |
| siwb_rough | 0.49 | -0.31 | -0.37 | -0.21 | 0.55 | 0.41 | -0.27 | 0.54 | 0.45 | 0.54 | 0.46 | 0.43 | 0.42 | -0.24 | 0.40 | 0.42 | -0.04 | 0.41 | -0.22 | 0.19 | -0.51 | 0.05 | 1.00 |  |  |  |
| siwb_slope | 0.01 | 0.16 | 0.08 | 0.20 | 0.19 | -0.07 | 0.17 | 0.09 | -0.03 | 0.09 | -0.03 | -0.08 | -0.07 | -0.20 | -0.14 | -0.08 | -0.18 | -0.09 | -0.13 | -0.05 | 0.21 | -0.06 | 0.00 | 1.00 |  |  |
| siwb_hills | -0.15 | 0.11 | 0.22 | 0.05 | -0.15 | -0.12 | 0.08 | -0.16 | -0.13 | -0.17 | -0.13 | -0.16 | -0.15 | 0.21 | -0.09 | -0.15 | 0.10 | -0.13 | 0.13 | -0.36 | 0.09 | -0.23 | -0.18 | -0.22 | 1.00 |  |
| siwb_aspec | -0.33 | 0.38 | 0.27 | 0.41 | -0.19 | -0.38 | 0.40 | -0.27 | -0.36 | -0.27 | -0.36 | -0.37 | -0.37 | -0.20 | -0.36 | -0.37 | -0.33 | -0.37 | -0.16 | -0.12 | 0.08 | -0.02 | -0.31 | 0.07 | 0.35 | 1.00 |
